# Lipids of different phytoplankton groups differ in sensitivity to degradation: implications for carbon export

**DOI:** 10.1101/2023.05.10.539996

**Authors:** Jelena Godrijan, Daniela Marić Pfannkuchen, Tamara Djakovac, Sanja Frka, Blaženka Gašparović

## Abstract

The future of life on Earth depends on how the ocean might change, as it plays an important role in mitigating the effects of global warming. The main role is played by phytoplankton. Not only are phytoplankton the base of the oceans’ food web, but they also play an important role in the biological carbon pump (BCP), the process of forming organic matter (OM) and transporting it to the deep sea, representing a sink of atmospheric CO_2_. Lipids are considered important vectors for carbon sequestration. A change in the phytoplankton community composition as a result of ocean warming is expected to affect the BCP. Many predictions indicate a dominance of small at the expense of large phytoplankton. To gain insight into interplay between the phytoplankton community structure, lipid production and degradation and adverse environmental conditions, we analyzed phytoplankton composition, POC and its lipid fraction in the northern Adriatic over a period from winter to summer at seven stations with a gradient of trophic conditions. We found that at high salinity and low nutrient content, where nanophytoplankton prevailed over diatoms, the newly fixed carbon is substantially directed toward the synthesis of lipids. Lipids produced by nanophytoplankton, coccolithophores and phytoflagellates, are more resistant to degradation than those produced by diatoms. This suggests a more successful lipid carbon sink of nanophytoplankton and thus a negative feedback on global warming. The difference in lipid degradability is discussed as a difference in the size of the cell phycosphere. We hypothesize that the lipids of nanophytoplankton are less degradable due to the small phycosphere with a poorer bacterial community and consequently a lower lipid degradation rate compared to diatoms. The chemical composition of the lipids of the different phytoplankton groups could have a different susceptibility to degradation, which could also contribute to the differences in lipid degradability.

## 1 INTRODUCTION

The ocean is important for the global carbon budget (Friedlingstein et al., 2022). It regulates atmospheric CO_2_ concentrations and is estimated to absorb 25% of annual anthropogenic carbon emissions (Heinze et al., 2015). The ocean carbon budget consists of inorganic and organic pools distributed between the particulate and dissolved fraction. The organic pool originates primarily from autochthonous sources and secondarily allochthonous sources (Lønborg et al., 2020). Autochthonous organic matter (OM) is produced by phytoplankton through photosynthesis from dissolved CO_2_ in a process known as primary production (PP). The produced OM is transferred downward through the ocean by the action of the biological carbon pump (BCP), mediated by either biological or physical processes (Claustre et al., 2021). The BCP sequesters carbon for weeks to hundreds or even millions of years (DeVries et al., 2012). How efficiently the BCP sequesters carbon at ocean depth depends in large part on the fraction of primary production exported below the euphotic zone (Buesseler and Boyd, 2009). The major factors determining BCP efficiency are particulate OM flux, net PP, food web controls, ballast, temperature, oxygen content, and degradation rates (Buesseler et al., 2020). Most of the OM is already removed or partially degraded in the surface layers of the ocean by the action of bacteria (Azam et al., 1983). However, it is important to emphasize that the lability of OM is one of the key factors determining the residence time of OM in the ocean (Cabrera-Brufau et al., 2021, Moran et al., 2021).

As a result of global change, the oceans’ PP is declining (2.1% decline per decade) (Gregg and Rousseaux, 2019). Phytoplankton play a key role in global PP, major biogeochemical cycles, and form the basis of the food chain in aquatic environments. The succession of dominant life-forms in phytoplankton is shaped by a complex interplay of many factors, including nutrient and light availability, temperature, and turbulence (Barbosa et al., 2010). Among the major phytoplankton groups, coccolithophores and diatoms with calcified and silicified cell walls, respectively, have global ecological significance, including the role they play in the global carbon cycle through the production and export of inorganic and organic carbon to the ocean depths (O’Brien et al., 2013; Gregg and Rousseaux, 2019). Global change is affecting phytoplankton biomass, primary productivity, and carbon export. It is evident that diatom abundance has declined significantly in many regions of the world’s oceans (Mishra et al., 2022), while the abundance of coccolithophores in the North Atlantic has increased in the past 50 years (Rivero-Calle et al., 2015). Observed changes in phytoplankton, including abundance (Boyce et al., 2010) and community structure (Marinov et al., 2010), are expected to have a cascading influence on primary and export production, food web dynamics, and marine food web structure (Chust et al., 2014).

Phytoplankton are the most important source of biogenic lipids in the ocean (Gašparović et al., 2014). The content and composition of biosynthesized lipids depend on environmental factors (Guschina and Harwood, 2009). Lipids are rich in carbon and are one of the major biochemicals in the ocean. Lipids with saturated acyl chain are shown to be selectively preserved in the water column, making them an important vector for carbon sequestration and potentially important factors in the efficiency of the BCP (Gašparović et al., 2016). Early diagenetic changes affect the chemical stability of lipids and their longevity in the water column (Brassell, 1993). Nonselective preservation of lipids could be enabled by physical protection through their association with minerals, such as diatom’s siliceous frustules and calcite coccoliths of coccolithophores (Hedges et al., 2001). In the water column, lipids are subjected to biotic (enzymatic peroxidation, biohydrogenation (Rontani and Koblížek, 2008)), and abiotic (photooxidation, autoxidation) (Rontani, 2008) breakdown processes. While autoxidation and biotransformation may take place throughout the water column, photooxidation may play a significant role in the euphotic layer (Rontani et al., 2009). While abiotic degradation predominated in the suspended particle pool, biotic (heterotrophic) degradation was significant for sinking particles and increased with depth (Christodoulou et al. 2009).

Influence of global warming on the ocean is not only seen through the increase in its temperature, but also through a number of indirect changes including: oligotrophication of the upper water column due to increased ocean stratification that reduces water column mixing, reduced CO_2_ solubility, ocean acidification, deoxygenation, and a reduction in thermohaline circulation (IPCC, 2021). Under increasingly nutrient-depleted conditions, smaller phytoplankton is favored at the expense of larger diatoms (Bopp et al., 2005). To gain insight into the interplay between phytoplankton structure, biogenic lipid production and degradation, and environmental conditions, we analyzed phytoplankton and lipid production in the northern Adriatic along the transect of a well-defined trophic gradient from winter to summer. We hypothesized that lipids of different phytoplankton groups differ in their susceptibility to degradation. We also hypothesize that lipids that are more resistant to degradation may contribute positively to the BCP.

## 2 MATERIALS AND METHODS

### 2.1 Sampling and parameter analyses

Data were collected on seven cruises on a monthly basis from February to August in 2010. Seven stations were sampled throughout the northern Adriatic, from the transect between Rovinj and the Po River delta, covering hydrodynamically and trophically distinct regions (Figure 1). Water samples were collected with 5 L Niskin bottles at the surface (0.5 m depth).

**FIGURE 1.**
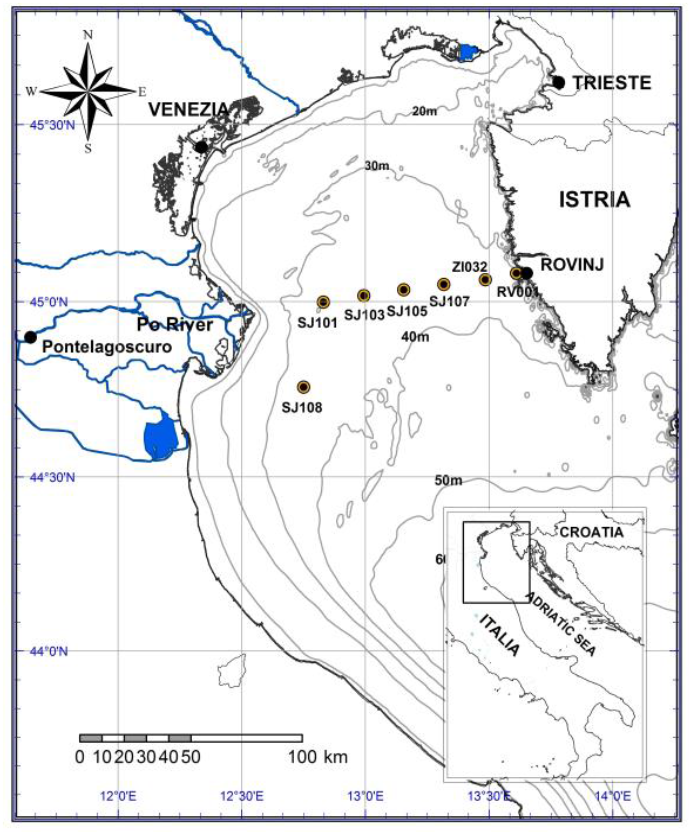
Map of sampling stations in the northern Adriatic Sea.

A CTD probe (Seabird SBE25, Sea–Bird Electronics Inc., Bellevue, Washington, USA) was used to measure temperature and salinity. Total phosphorus, dissolved inorganic orthophosphates (PO_4_^3−^), total inorganic nitrogen (TIN), including nitrates (NO_3_^−^), nitrites (NO_2_^−^), and ammonium (NH_4_^+^)), were determined by spectrophotometric methods (Parsons et al., 1984) on board and immediately after sampling using Shimadzu UV-Mini 1240 spectrophotometer with 10 cm quartz cuvettes. Organic phosphorus concentration was calculated as the difference between total and inorganic phosphorus concentrations. Subsamples for the determination of chlorophyll *a* (Chl *a*) were filtered on Whatman GF/C filters and stored frozen at −20°C until further processing. Chl *a* concentrations were determined following 3 h extraction in 90% acetone (in the dark, with grinding), on a Turner TD–700 fluorimeter (Parsons et al., 1984).

### 2.2 Phytoplankton

We preserved 200 mL of seawater with 2% neutralized formaldehyde (final concentration) and performed nano- and microphytoplankton determination and enumeration within one month of sampling. The stored sample was homogenized by gentle shaking, and a subsample was added to the Utermöhl sedimentation chamber (volume: 50 mL; Hydro-Bios Apparatebau, Altenholz, Germany), where it settled for ∼30 h. We performed the analysis on a Zeiss Axiovert 200 (Zeiss, Jena, Germany) following the inverted microscope method (Utermöhl, 1958, Hasle 1978). Total phytoplankton included all species counted in the microphytoplankton (20–200 μm) and nanophytoplankton (2-20 μm) groups (Sieburth et al., 1978). Identified taxa were grouped to diatoms, dinoflagellates, and nanophytoplankton coccolithophores and phytoflagellates (which included chlorophytes, chrysophytes, cryptophytes and prasinophytes) according to Tomas (1997).

### 2.3 Particulate organic carbon (POC)

For POC determination, 1 L of seawater was filtered on board through 0.7 μm Whatman GF/F filters precombusted at 450 °C/5h. After filtration, the filters were rinsed with Milli-Q water to remove salts and stored in liquid nitrogen on board and at −80 °C in the laboratory until analysis. POC was analyzed using an SSM–5000A solid sample module connected to a Shimadzu TOC– V_CPH_ carbon analyzer calibrated with glucose (Sugimura and Suzuki, 1988). POC concentrations were corrected based on filter blank measurements. The average filter blank value including the instrument blank value corresponded to 5 μg C L^−1^. The reproducibility obtained for the glucose standard was 3%.

### 2.4 Lipids

For particulate lipid analysis, we collected 3 L of seawater prefiltered through a 200 μm stainless steel screen to remove larger particles including microzooplankton. Lipids were collected on through precombusted (450 °C/5h) 47 mm GF/F filters and stored in liquid nitrogen until lipid extraction. It was performed using a modified one-phase solvent mixture of dichloromethane-methanol-water procedure (Bligh and Dyer, 1959; Vrana et al., 2022). In short, in order to assess recoveries in later stages of sample analysis we added 5 μg of standard methyl stearate to the sliced filters together with 10 mL of a one-phase solvent mixture (dichloromethane/methanol/deionized water (1:2:0.8 v/v/v)). This was then subjected to an ultrasonic treatment for three minutes and stored overnight in the refrigerator, afterwards we filtered the extracts through a sinter funnel into a separatory funnel, washed once with a the one-phase solvent mixture, once with dichloromethane and 0.73% NaCl (1:1 v/v), and once with dichloromethane. The extracts were concentrated by rotary evaporation under a nitrogen atmosphere and kept at −20 °C until measurements were made. To prepare the lipid extracts for analysis, the dichloromethane extracts were evaporated to dryness under nitrogen flow and then dissolved in 20 μL dichloromethane prior to analysis.

Lipid classes were separated on Chromarods SIII and quantified with external calibration using a mixture of standard lipids by a thin-layer chromatograph-flame ionization detector (TLC-FID) Iatroscan Mark-VI (Iatron), using a hydrogen flow of 160 mL min^−1^ and an air flow of 2000 mL min^−1^. This method identify eighteen lipid classes: hydrocarbons (HC), steryl esters (SE), fatty acid methyl esters (ME), fatty ketone (KET), triacylglycerols (TG), free fatty acids (FFA), fatty alcohols (ALC), 1,3-diacylglycerols (1,3DG), sterols (ST), 1,2-diacylglycerols (1,2DG), pigments (PIG), monoacylglycerols (MG), three glycolipids (GL) including monogalactosyl-, digalactosyl-, and sulfoquinovosyl-diacylglycerol (MGDG, DGDG, and SQDG, respectively), and three phospholipids (PL) (phosphatidylglycerols (PG), phosphatidylethanolamines (PE), and phosphatidylcholines (PC)). Total lipid concentration is calculated by summing all detected classes. Full details can be found in Gašparović et al. (2015; 2017). In this article we focused on lipid degradation indices trough the lipolysis index (Goutx et al., 2003), which characterize the degree of lipid degradation in seawater. Lipolysis index is calculated as the ratio of the sum of lipid degradation indices (ALC+FFA+MG+DG) to the sum of cell lipids TG, WE, and glyco- and phospho-lipids (Goutx et al., 2003).

### 2.5 Data analysis

Linear and polynomial fits (Origin 7 computer software, Origin Lab) were performed to examine the correlation between salinity and major nutrient distributions, major phytoplankton groups developed, and lipid production in the northern Adriatic Sea.

To investigate correlations between major phytoplankton groups developed under different environmental conditions (salinity (S), temperature (T), major nutrients) and total organic matter (POC) and lipid production Principal component analysis was carried out using Statistica software. Schematic representation was drawn using symbols courtesy of the Integration and Application Network, University of Maryland Center for Environmental Science (ian.umces.edu/symbols/).

## 3 RESULTS

### 3.1 Environmental conditions

Sea surface salinity increased from stations in the Po River plume influence area on the western side of the northern Adriatic (stations SJ108, SJ101, SJ103) to the eastern side (stations SJ105, SJ107, ZI032 and RV001) (Figure 2a). The variability of sea surface temperature and salinity showed similar patterns. From winter to summer, water freshening occurred in parallel with the temperature increase (Figure 2b). Exceptions were observed during the March, May, and June at stations SJ108, SJ101, and SJ103, when colder river water mixed with warmer seawater. The decrease in salinity resulted in a substantial increase in inorganic nutrients, TIN (Figure 2c) and PO_4_^3−^ (Figure 2d), along with an increased ratio of TIN and total phosphorus (TIN/P_tot_) (Figure 2f). Total phosphorus rather than PO_4_^3−^ was used for the calculation of TIN/P_tot_ because organic phosphorus (Figure 2e) is an important source of phosphorus for phytoplankton in the northern Adriatic (Ivančić et al., 2012). The data used to create Figure 2 can be found in Table S1.

**FIGURE 2.**
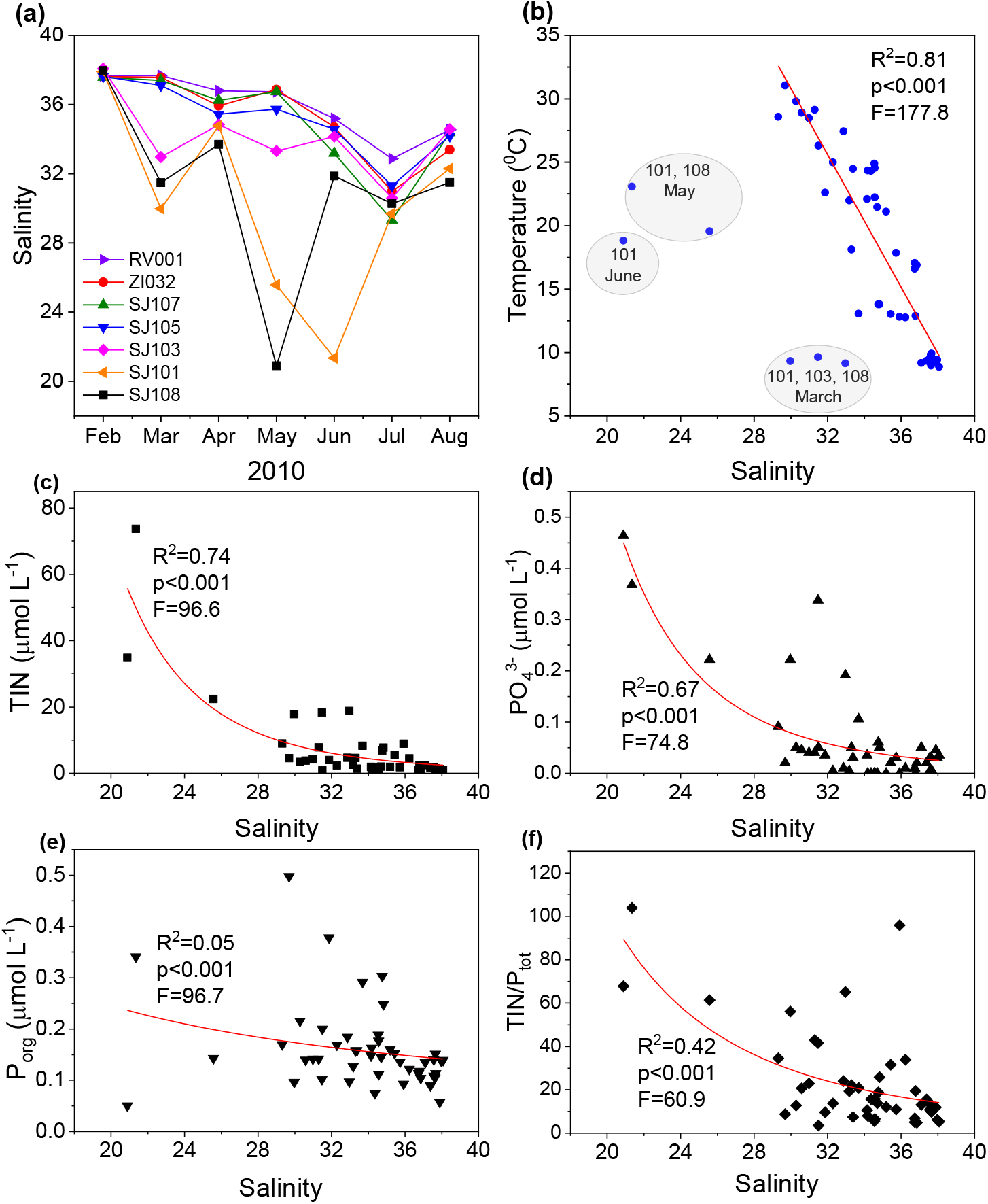
Relationship between salinity and other environmental parameters. Temporal salinity variations (A), relationships between salinity and temperature (B), TIN concentration (C), PO_4_^3−^ (D), organic phosphorus (E) concentrations and TIN/P_tot_ ratio (F) in the period from February to August 2010 at seven stations in the northern Adriatic. Note that linear fit in (b) is made without outliers separated by circles.

### 3.2 Phytoplankton succession

Phytoplankton community structure varied with salinity and time (Figures 3 and 4, Figure S1, Table S1), indirectly indicating a dependence of different phytoplankton groups on nutrient availability. Diatoms thrived at the lowest observed salinities (Figure 3a) and were associated with an increase in temperature (Figure 4). Coccolithophores thrived under conditions of high salinity (Figures 3b and 4) characterized by low temperatures (Figure 2b) and low nutrient availability (Figures 2d-e). On the spatio-temporal scale, phytoflagellates in the nano fraction found their niche for development at intermediate salinities (around 31-37), mainly at oligotrophic stations and from February to August (Figures 3c and 4, Figure S1). Dinoflagellates were abundant mainly in May and June (Figure 4, Figure S1) at intermediate salinities 30-36.5 (Figure 3d). The temporal succession of dominant phytoplankton groups is shown in Table 1.

**FIGURE 3.**
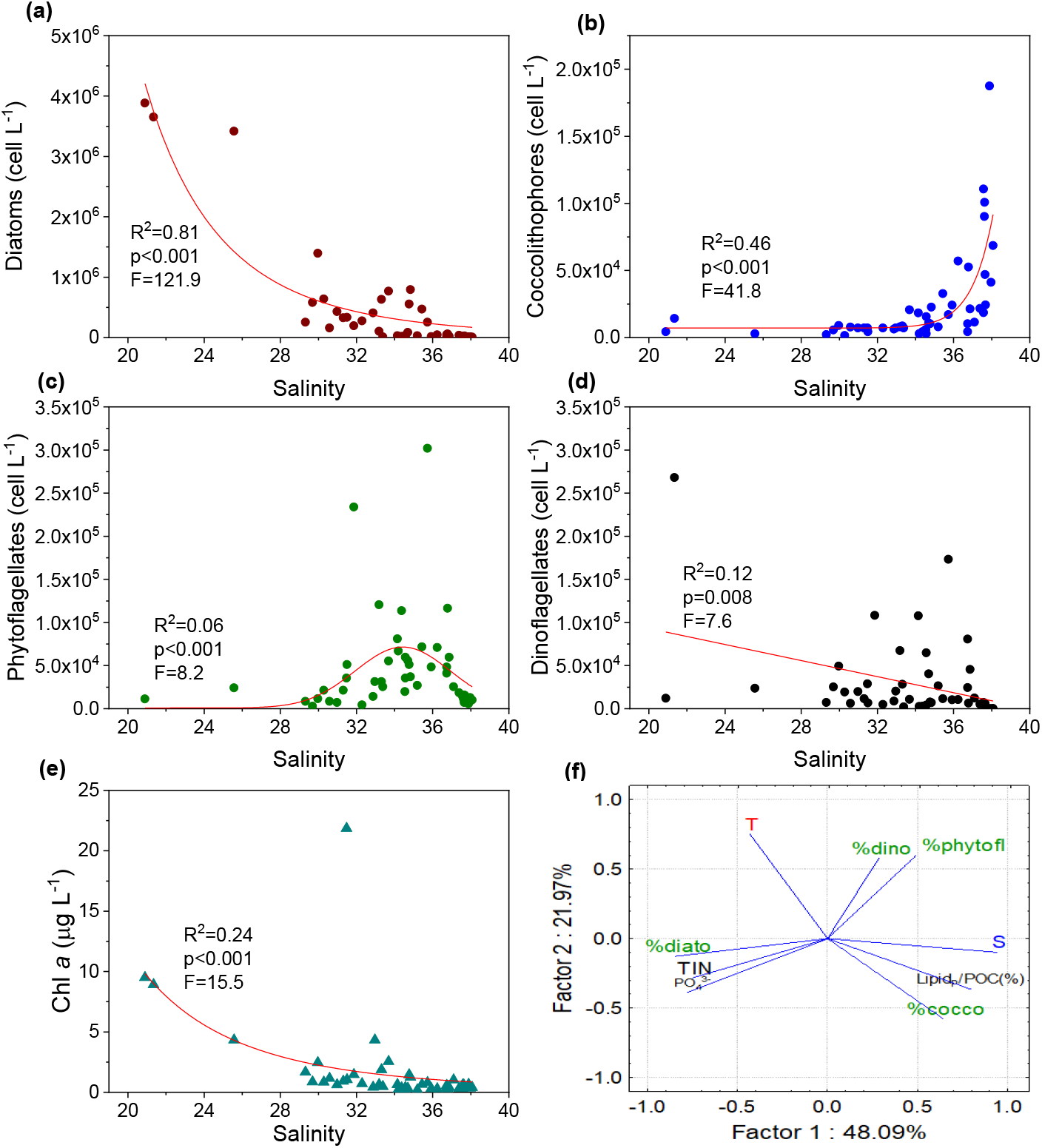
Relationship between salinity and abundance of micro diatoms (A), coccolithophores (B), phytoflagellates (C), dinoflagellates (D), Chl *a* (E). Biplot of scores of the contribution of major phytoplankton groups (%, diatoms, coccolithophores, phytoflagellates, and dinoflagellates), the relative content of particulate lipids in POC (Lipid_P_/POC (%)) and environmental factors (T - temperature, S - salinity, TIN - total inorganic nitrogen and PO_4_^3−^ - ortophosphates) from the results of principal component analysis (F).

**FIGURE 4.**
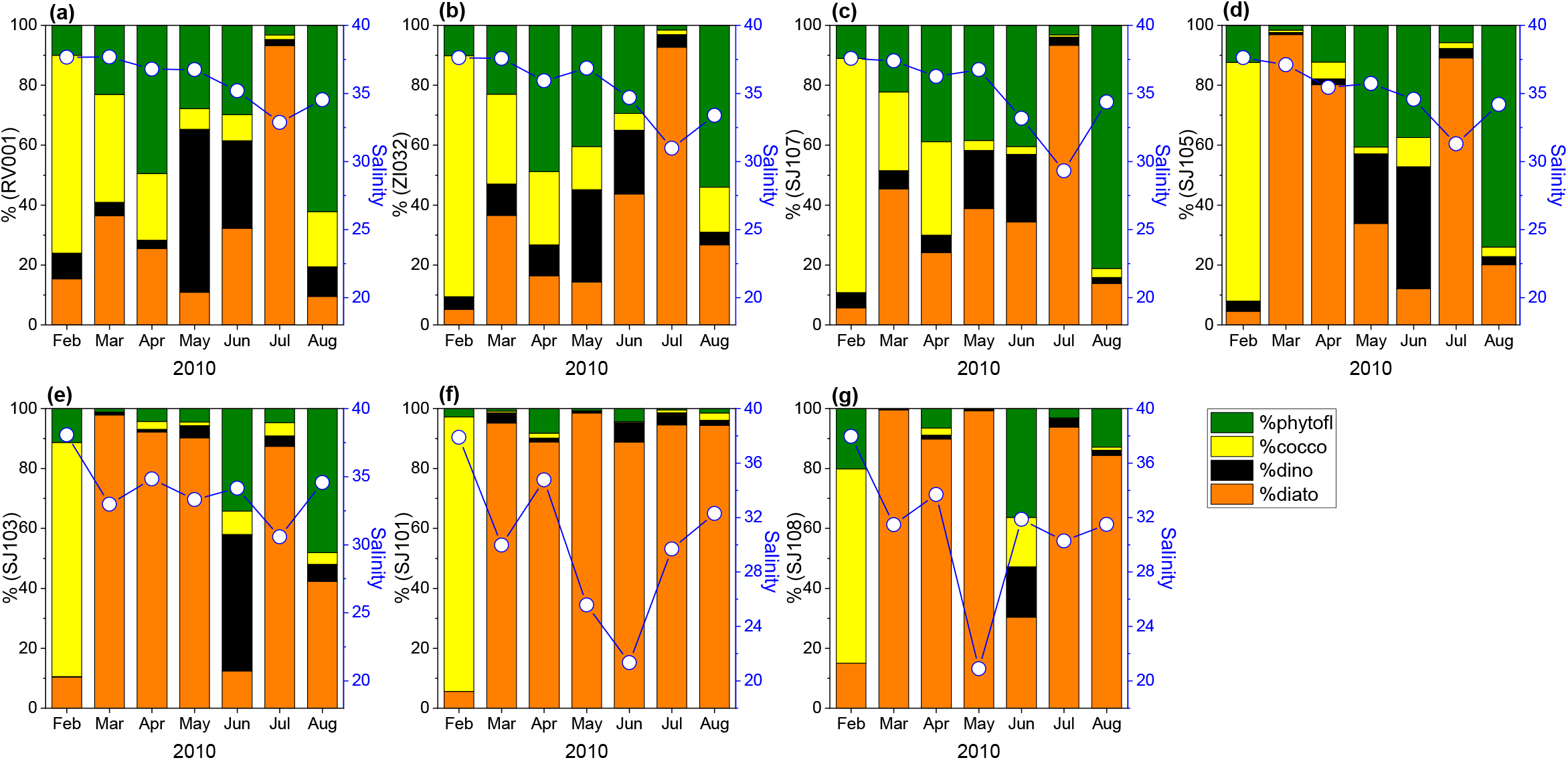
Phytoplankton community structure (bars, % of total community obtained by microscopic identification and counting of the cells) for the period February to August in the surface waters (0.5 m depth) of the northern Adriatic Sea at stations RV001 (a), ZI032 (b), SJ107 (c), SJ105 (d), SJ103 (e), SJ101 (f) and SJ108 (g). Temporal salinity variations (right y-axis) are shown for each station with lines and symbols.

**Table 1.** Temporal succession of the dominant phytoplankton group(s) and environmental data (salinity and Chl *a*) for the seven cruises conducted in the northern Adriatic Sea from February to August 2010.

| Date | Stations | Dominant phytoplankton group(s) | Chl *a*±SD<br>(µg/L) | Average<br>salinity±SD |
| --- | --- | --- | --- | --- |
| 15.02.2010 | All | Coccolithophores | 0.51±0.12 | 37.77±0.20 |
| 17.03. 2010 | RV001-SJ107 | Mixed, mainly nanophytoplankton<br>with significant proportion of<br>coccolithophores (26.2–35.9%) | 0.36±0.05 | 37.55±0.15 |
|  | SJ105-SJ108 | Diatoms | 2.61±9.72 | 33.35±3.07 |
| 15.04. 2010 | RV001-SJ107 | Mixed, mainly nanophytoplankton<br>with significant proportion of<br>coccolithophores (22.3–31.2%) | 0.30±0.02 | 36.32±0.44 |
|  | SJ105-SJ108 | Diatoms | 1.48±0.79 | 34.68±0.73 |
| 17.05. 2010 | RV001-SJ105 | Mixed, mainly nanophytoplankton | 0.54±0.20 | 36.52±0.53 |
|  | SJ103-SJ108 | Diatoms | 5.24±3.90 | 26.59±6.27 |
| 24.06. 2010 | SJ108-RV001 | Mixed, mainly nanophytoplankton | 1.80±3.16 | 33.94±1.22 |
|  | SJ101 | Diatoms | 5.19±5.26 | 21.35±0.00 |
| 15.07. 2010 | All | Diatoms | 0.93±0.66 | 30.72±1.78 |
| 19.08. 2010 | RV001-SJ103 | Mixed, mainly nanophytoplankton | 0.29±0.15 | 34.21±0.63 |
|  | SJ101 and SJ108 | Diatoms | 0.88±0.23 | 31.90±0.56 |

In the PCA analysis (Figure 3f, Table S2), PC1 and PC2 explained 70.06% of the variances and revealed positive relationship between high salinity and increase in coccolithophore proportion to the total phytoplankton (%cocco) and the relative content of particulate lipids in POC (Lipid_P_/POC (%)). On the other hand, the contribution of diatoms to the total phytoplankton (%diato) is clustered with the inorganic nutrients, TIN and PO_4_^3−^. PC1 clearly distinguishes these two clusters.

### 3.3 Lipid production and degradation

In parallel with Chl *a* (Figure 3e), high OM content characterized low salinity samples, including POC (Figure 5a) and particulate lipid content (Figure 5b). Conversely, the relative content of particulate lipids in POC (Lipid_P_/POC (%)) increased significantly in high salinity waters (Figure 5c), characterized by low POC (Figure 5a) and high percentage of coccolithophores (Figure 3b). The data used to create Figure 5 can be found in Table S1.

**FIGURE 5.**
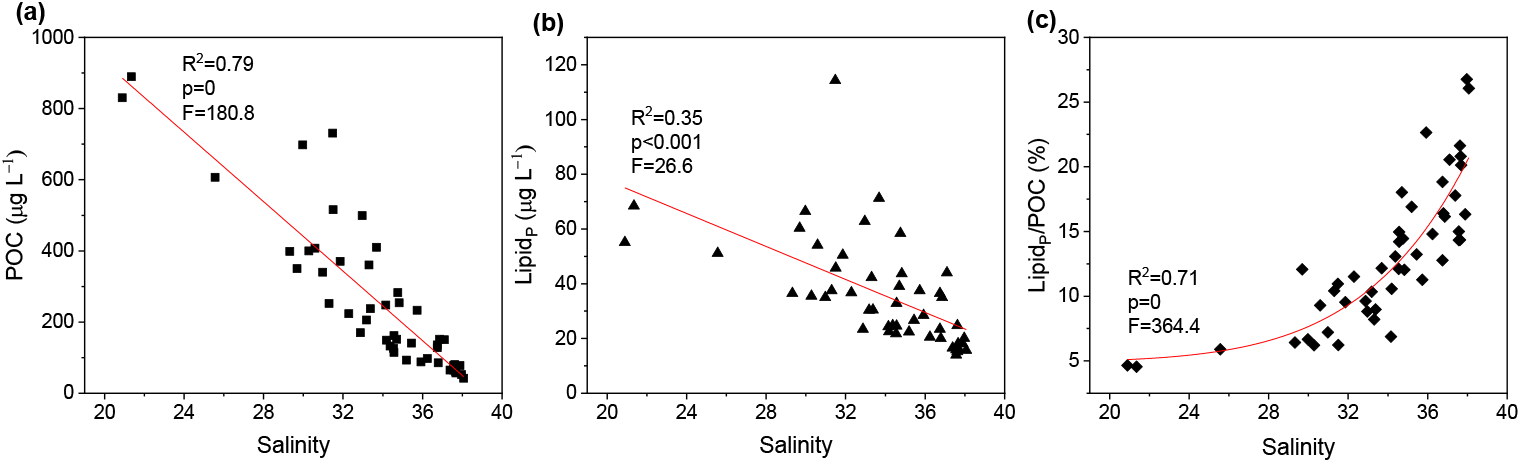
Relationships between salinity and POC (a), particulate lipid concentration (b), the relative content of particulate lipids in POC (c).

For the stations and time periods where the total population consisted of coccolithophores or a mixed population, composed mainly of nanophytoplankton, a much higher lipid content per Chl *a* was observed, 157–417 pg lipid/Chl *a* than in the cases where the total population was dominated by diatoms, 21–120 pg lipid/Chl *a* (Figure 6a). The correlation of Lipid_P_/POC (%) with major phytoplankton groups indicated smaller Lipid_P_/POC (%) when diatoms were abundant and, conversely, higher lipid_P_/POC (%) when coccolithophores were abundant.

**FIGURE 6.**
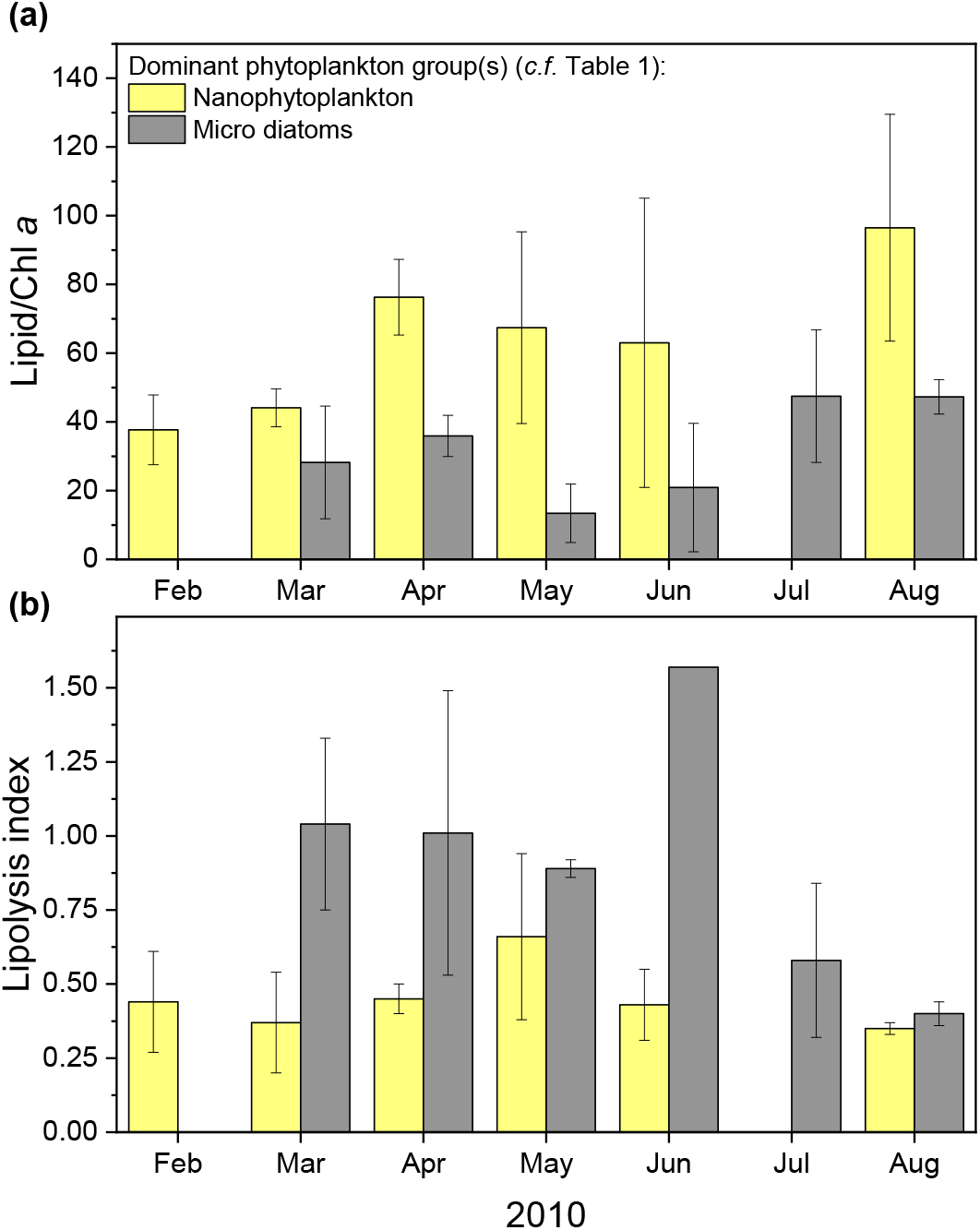
Lipid production (Lipid/Chl *a*) and degradation (Lipolysis index (LI)) in the period from February to August 2010 in the northern Adriatic Sea.

The total particulate lipid fraction analyzed also contained lipids from microzooplankton, whose carbon yield in the northern Adriatic POC is orders of magnitude lower than that of phytoplankton (Kamburska and Fonda-Umani, 2009) as well as from the bacterial community, but their contribution to total lipids is considered to be neglected. The contribution of bacterial carbon to POC is generally low, ranging from 1-17% in the northern Adriatic (La Ferla et al., 2005) and 1-2% in the North Atlantic (Gašparović et al., 2014). At the same time, the lipid carbon of marine bacteria is low and ranges from 1.7 to 7.3% of the POC (Goutx et al., 1990). Lipids may have also partially originated from the non-living OM, which could have contributed significantly to the POC and lipid pool of the phytoplankton growing under stress conditions (Flanjak et al., 2022).

An index to characterize the degree of lipid degradation in natural seawater and OM freshness is the lipolysis index (LI) (Goutx et al., 2003). Higher LI values indicate a higher degree of lipid degradation. Higher LI values were determined for stations and time periods characterized by the dominance of diatoms (LI=0.40–1.57) with respect to the dominance of coccolithophores or mixed, mainly nanophytoplankton population (LI=0.35–0.66) (Figure 6b).

## 4 DISCUSSION

### 4.1 Different sensitivity to decomposition of lipids of different phytoplankton groups

We found that under conditions of high salinity and low inorganic nutrient content (oligotrophic area), nanophytoplankton prevailed over microphytoplankton, and the newly fixed carbon appeared to be substantially directed toward the synthesis of lipids, as indicated by the high relative content of particulate lipids in POC (> 15%, Figure 5c) and the high lipid content per Chl *a* (Figure 6a). In contrast, excess nutrients at low salinity (mesotrophic area) promoted diatom blooms (Figure 3a), which synthesized more lipids, but the relative content of particulate lipids in POC (< 5%) and the lipid content per Chl *a* (Table 1) are low. These results suggest that diatom cell growing in nutrient abundant conditions have a low requirement for lipids as structural components of membranes, signal transduction, and energy storage.

Given that the total amount of lipids is higher during diatom blooms, the question is whether this necessarily means that diatoms (rich in OM) are more significant vectors of lipid carbon removal to the ocean depths in comparison to nanophytoplankton? We can also ask whether lipids produced by nanophytoplankton, coccolithophores and phytoflagellates, and diatoms have a different tendency to degrade. In addition to the many factors that influence the efficiency of transfer of OM to the depth, lower degradation likely favors longer residence time in the water column and most likely better efficiency of carbon sequestration.

The lower values of Lipolysis index under conditions in which coccolithophores pervaded or contributed significantly to the total population (Figure 6b) indicate that lipids produced by nanophytoplankton are more resistant to degradation than those produced by diatoms. This suggests that a greater proportion of nanophytoplankton lipids can be transported to depth than those produced by diatoms. One explanation for the difference in lipid stability between these phytoplankton groups could be the size of the cell phycosphere harboring a variety of bacterial species (Seymour et al., 2017). The size of the phycosphere is closely related to the phytoplankton cell size with the phycosphere radius of diatoms being one to five orders of magnitude larger than that of nanophytoplankton (Seymour et al., 2017), suggesting a much richer bacterial community hosted by the phycosphere of diatoms and, accordingly, more intense lipid degradation upon cell death.

The difference in lipid stability between phytoplankton groups could be also due to difference in lipid chemical composition, which affects the different susceptibility to degradation. Saturated lipids are more resistant to degradation and are therefore important carbon carriers to marine depths (Gašparović et al., 2016). Accordingly, abyssal depths of the Atlantic are enriched in saturated fatty acids, saturated or monounsaturated triacylglycerols (Gašparović et al., 2016). Coastal phytoplankton populations growing under N and P limitation biosynthesize saturated fatty acids, especially under P limitation (Grosse et al., 2019). Shin et al. (2003) found higher production of unsaturated fatty acids (fivefold) at the East China Sea mesotrophic site, which is characterized by the dominance of diatoms. In contrast, saturated fatty acids dominated at the oligotrophic sites, while the proportion of diatoms decreased.

Mayzaud et al. (2014) found that in the northern Atlantic, the proportion of polyunsaturated fatty acids was much greater under mesotrophic conditions than under oligotrophic conditions. Because diatoms from our study grew under higher/er nutrient availability, i.e. lower salinity (Figure 3a) than nanophytoplankton we can assume that diatoms synthesized more unsaturated lipids that are more susceptible to degradation (abiotic photodegradation or biotic). Consequently, we found higher values of lipolysis index were estimated for samples in which diatoms predominated. Furthermore, haptophytes, especially the coccolithophorid family Gephyrocapsaceae, are rich in long-chain (C37–C39) unsaturated ethyl and methyl ketones (Sawada and Shiraiwa, 2004). These long-chain alkenones are paleomarkers for reconstructing past changes in sea surface temperatures and global climate (Brassell et al., 1986, Volkman et al., 1995) because they are well preserved in bottom sediments and have also been used to reconstruct the paleoproductivity of coccolithophores (Schulte et al., 1999). This further suggests their stability to degradation and the importance of coccolithophore lipid carbon for carbon sequestration in the ocean (Raja and Rosell-Melé, 2021).

We also considered whether higher temperature (with linked higher bacterial activity) is the reason for enhanced diatom lipid degradation. For example, in March, when the temperature throughout the transect was 9.35 ± 0.18 °C (Table S1), the lipolysis index was low (average 0.45) at three oligotrophic stations (RV001, ZI032, and SJ107) with a significant presence of coccolithophores, indicating lower lipid degradation. In contrast, at the mesotrophic stations SJ105-SJ108, characterized by the dominance of diatoms, significantly higher lipid degradation was observed (average LI = 1.04), suggesting more intense diatom lipid degradation. Thus, the temperature effect in different degradation of lipids produced by particular phytoplankton groups could be neglected (Figure 6b).

### 4.2 Perspectives in a context of global climate change

If the changes in phytoplankton community structure predicted by Bopp et al. (2005) and Gregg and Rousseaux (2019) occur in the context of climate change, lipids from coccolithophores and other nanophytoplankton could play a significant role in the process of carbon sequestration to the deep sea (Figure 6). Natural populations dominated by small flagellates and coccolithophores are often rich in lipids, as studies have shown both in model experiments (Fernández et al. 1994a, up to 60% of C in lipids) and in open ocean waters (Fernández et al., 1994b), consistent with our results. In contrast, diatoms growing under favorable nutrient and temperature conditions do not accumulate lipids (e.g., Flanjak et al., 2022), which is also consistent with our results. However, warming and lack of nutrients affect the increased incorporation of carbon into lipids in diatom cells (e.g. Novak et al., 2019). The coccolithophore *Emiliania huxleyi* accumulates much more lipids in N-poor conditions (69.0 pg/cell) compared to N-replete growth conditions (11.4 pg/cell) (Pantorno et al., 2013).

Warming generally affects reduced polyunsaturated fatty acid biosynthesis (Hixson and Arts, 2016). In diatoms, an increase in temperature leads to fatty acid shortening and a decrease in total unsaturated fatty acid content in both galactolipids (Dodson et al., 2014) and phospholipids (Vrana et al., 2022). Whereas, warming leads to a decrease in the carbon-normalised content of PUFAs in the coccolithophore *E. huxleyi* (Bi et al., 2020). Lower lipid saturation would result in lower susceptibility to degradation, resulting in better export of lipid carbon to the ocean interior. The lower lipid degradability has a positive effect on BCP, as its efficacy is also determined by the rate of OM/lipid degradation (Buesseler et al., 2020).

Lipids are generally discussed to be buoyant and as such do not settle but remain in the water column. However, due to their surface-active properties, they escape from the water and attach to the sinking particles, thus contributing to carbon sequestration (Novak et al., 2018).

Other predictions also suggest a positive effect of coccolithophores on BCP compared to diatoms. Even though the modelling study of the distribution of the diatom *Chaetoceros diadema* and the coccolithophore *E. huxleyi* in the ocean predicts their future decline (Jensen et al., 2017), they also note that the probability of *C. diadema* settling below 1000 m would decrease significantly, while the abundance of *E. huxleyi* at these depths would not change significantly from current conditions. In addition, a recent study using new and historical data has shown that *E. huxleyi* was found not affected by high temperatures during ‘hot summers’ and its abundance increases as the warming trend continues, indicating its ability to thrive and adapt to ocean warming (Frada et al., 2021). Nutrient limiting conditions may have a positive effect on the sinking rate of coccolithophores. The coccolithophore *Gephyrocapsa oceanica* has a higher sinking rate associated with higher calcification for growth under N-limited conditions (Jiang et al., 2022). Moreover, *E. huxleyi* from the stationary growth phase that grew under N-limitation was also found to sink faster compared to growth under N-rich conditions (Pantorno et al., 2013). Lastly, Wang et al. (2022) reported that P-limitation might promote sinking of *E. huxleyi*.

In summary, our results suggest that diatoms (rich in OM) are probably not more significant vectors of lipid carbon removal to the deep ocean compared to nanophytoplankton. If nanophytoplankton predominate over diatoms in the future oceans, the carbon sink via lipids will be higher than expected based only on the lower carbon content in nanophytoplankton cells compared to diatoms (Figure 7) for at least two reasons: 1) nanophytoplankton synthesize relatively more lipids than larger diatoms, and 2) the lipids of nanophytoplankton are much less susceptible to degradation than those of diatoms and therefore can remain in the water column longer, be transferred deeper and potentially be stored in oceanic sediments. In such an environment, the lipid carbon may be preserved for millions of years.

**FIGURE 7.**
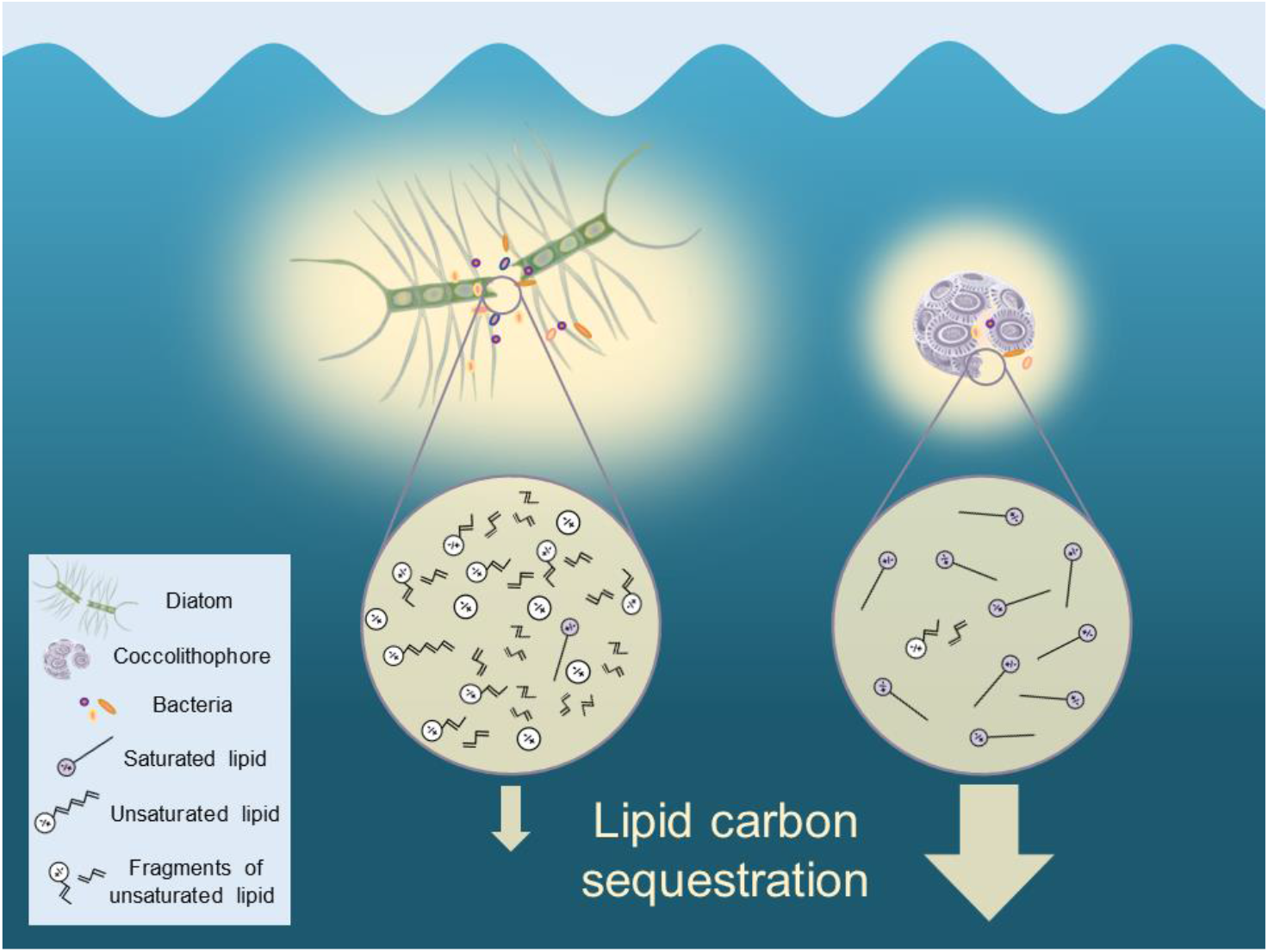
Schematic representation of the interplay between diatom and coccolithophore (representative of nanophytoplankton) phycospheres, lipid composition and degradability, and relationship to carbon sequestration efficiency.

## AUTHOR CONTRIBUTIONS

JG and BG conceived of and designed the study. JG, SF, DMP and TD conducted the experiments and analyzed the data. BG prepared the original draft of the manuscript with writing, reviewing, and editing from JG, SF, DMP and TD.

## ACKNOWLEDGMENTS

This work was funded by grants from the Croatian Science Foundation under projects UIP-2020-02-7868, IP-2018-01-3105 and IP-11-2013-8607 and the Croatian National Monitoring Program (Project “Jadran”). We thank Margareta Buterer, Jasna Jakovčević, Paolo Krelja and the crew of the RV “Vila Velebita” for their help in sampling.

## DATA AVAILABILITY STATEMENT

The data used to create Figures can be found in Table S1.

