## Supplementary material for "Lipids of different phytoplankton groups differ in sensitivity to degradation: implications for carbon export": Supplemental_Files_Godrijan_GCB_bioRxiv.pdf

### The lipids of different phytoplankton groups have different sensitivities to degradation: implication for carbon export in the future ocean

**Jelena Godrijan[1], Daniela Marić Pfannkuchen[2], Tamara Djakovac[2], Sanja Frka[1], Blaženka Gašparović[1]**

[1]Division for Marine and Environmental Research, Ruđer Bošković Institute, POB 180, HR–10002 Zagreb, Croatia
[2]Center for Marine Research (CMR), Ruđer Bošković Institute, G. Paliaga 5, 52210 Rovinj, Croatia

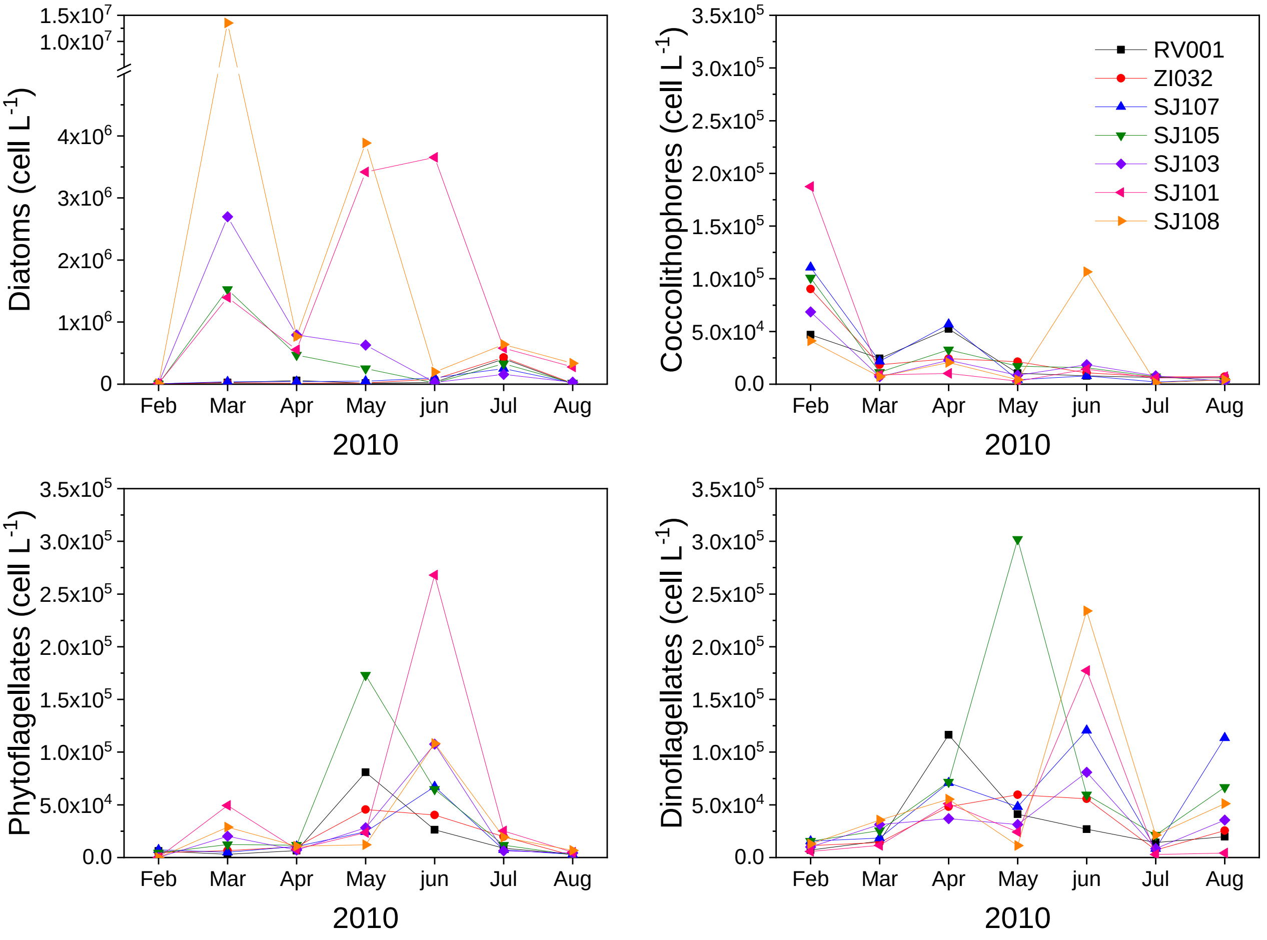


FIGURE S1 The abundance of the most important phytoplankton groups for the period February to August in the surface waters (0.5 m depth) of the investigated stations in the northern Adriatic Sea.

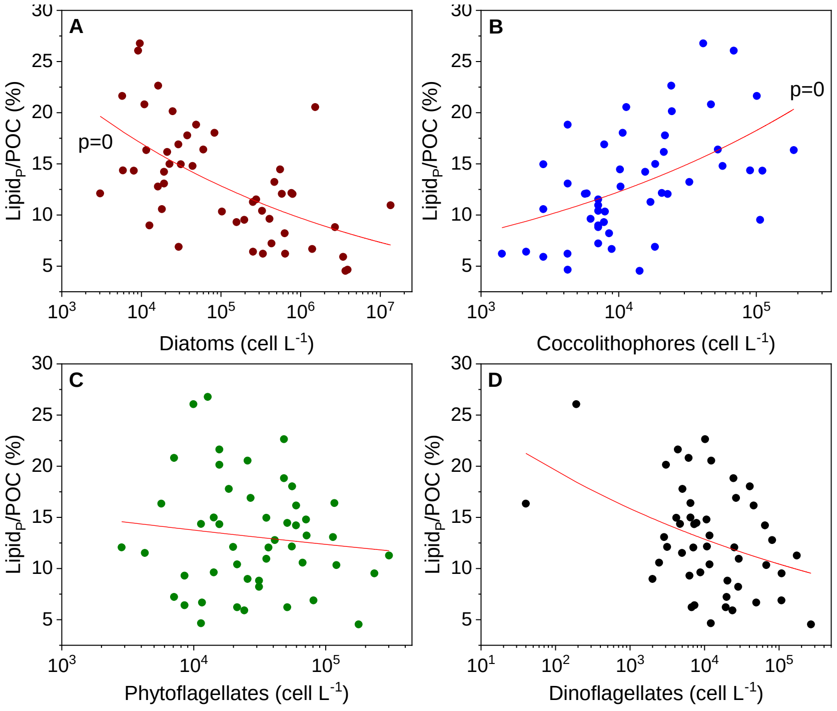


FIGURE S2 The relationship between relative particulate lipid abundance in POC (Lipid$_P$/POC (%)) and micro diatoms (A), coccolithophores (B), phytoflagellates (C), dinoflagellates (D) abundances.

TABLE S1 The data used to create Figures 2-6, and S1.

| Date | Station | S | T | $PO_4^{3-}$ | $P_{org}$ | TIN | Chl *a* | POC | $Lipid_P$ | DI* | Cell lipids | $Lipid_P$ | Lipolysis index | diato | cocco | phytofl | dino |
|---|---|---|---|---|---|---|---|---|---|---|---|---|---|---|---|---|---|
| | | | $^0$C | | µmol $L^{-1}$ | | | | µg $L^{-1}$ | | | % | | | cell/L | | |
| 15/02/10/ | RV001 | 37.66 | 9.91 | 0.00 | 0.15 | 1.49 | 0.46 | 60.87 | 18.1 | 5.0 | 10.8 | 20.8 | 0.46 | 2.4E+03 | 46860 | 7100 | 6100 |
| | ZI032 | 37.62 | 9.83 | 0.02 | 0.11 | 1.48 | 0.45 | 74.35 | 15.2 | 2.7 | 11.5 | 14.4 | 0.23 | 4.5E+03 | 90220 | 11360 | 4680 |
| | SJ107 | 37.57 | 9.55 | 0.02 | 0.14 | 1.57 | 0.60 | 77.41 | 15.9 | 3.3 | 11.0 | 14.3 | 0.30 | 2.4E+03 | 110760 | 15620 | 7260 |
| | SJ105 | 37.63 | 8.97 | 0.03 | 0.11 | 1.38 | 0.61 | 80.08 | 24.8 | 10.2 | 13.1 | 21.6 | 0.78 | 4.3E+03 | 100820 | 15620 | 4380 |
| | SJ103 | 38.07 | 8.86 | 0.04 | 0.14 | 0.93 | 0.41 | 42.21 | 15.7 | 4.8 | 10.1 | 26.1 | 0.48 | 6.3E+03 | 68540 | 9940 | 190 |
| | SJ101 | 37.89 | 9.26 | 0.05 | 0.06 | 1.23 | 0.66 | 77.54 | 18.1 | 4.8 | 11.1 | 16.3 | 0.43 | 8.7E+03 | 187440 | 5680 | 40 |
| | SJ108 | 37.97 | 9.42 | 0.03 | 0.14 | 1.05 | 0.35 | 52.5 | 20.1 | 4.8 | 12.3 | 26.8 | 0.39 | 6.0E+03 | 41180 | 12780 | 0 |
| | RV001 | 37.68 | 9.56 | 0.01 | 0.11 | 1.29 | 0.41 | 56.65 | 16.3 | 3.9 | 11.5 | 20.1 | 0.34 | 1.2E+04 | 24330 | 15620 | 3030 |
| 17/03/10/ | ZI032 | 37.57 | 9.24 | 0.03 | 0.11 | 1.88 | 0.33 | 65.18 | 14.0 | 1.7 | 8.2 | 15.0 | 0.21 | 1.3E+04 | 18460 | 14200 | 6480 |
| | SJ107 | 37.39 | 9.34 | 0.02 | 0.09 | 1.71 | 0.33 | 65.05 | 16.5 | 5.5 | 10.0 | 17.8 | 0.55 | 3.3E+04 | 21680 | 18460 | 5050 |
| | SJ105 | 37.10 | 9.18 | 0.05 | 0.14 | 2.43 | 1.02 | 150.1 | 44.1 | 17.8 | 23.2 | 20.5 | 0.77 | 1.4E+06 | 11360 | 25560 | 12280 |
| | SJ103 | 32.97 | 9.15 | 0.19 | 0.10 | 18.81 | 4.33 | 499 | 62.8 | 30.7 | 29.4 | 8.8 | 1.04 | 2.7E+06 | 7100 | 31240 | 20260 |
| | SJ101 | 29.97 | 9.32 | 0.22 | 0.10 | 17.86 | 2.46 | 697.4 | 66.5 | 29.2 | 32.2 | 6.7 | 0.91 | 1.4E+06 | 8900 | 11550 | 49430 |
| | SJ108 | 31.48 | 9.63 | 0.34 | 0.10 | 18.30 | 21.86 | 730.6 | 114.3 | 61.0 | 42.1 | 11.0 | 1.45 | 1.3E+07 | 7100 | 35500 | 28800 |
| 15/04/10/ | RV001 | 36.79 | 12.88 | 0.00 | 0.12 | 2.29 | 0.29 | 85.35 | 20.0 | 5.9 | 12.7 | 16.4 | 0.47 | 4.3E+04 | 52503 | 116358 | 6456 |
| | ZI032 | 35.92 | 12.81 | 0.00 | 0.09 | 8.90 | 0.32 | 87.89 | 28.4 | 7.6 | 18.3 | 22.6 | 0.42 | 1.5E+04 | 24123 | 48246 | 10175 |
| | SJ107 | 36.24 | 12.76 | 0.01 | 0.12 | 4.46 | 0.29 | 97.1 | 20.5 | 6.2 | 13.0 | 14.8 | 0.48 | 4.3E+04 | 56950 | 70950 | 10644 |
| | SJ105 | 35.44 | 13.03 | 0.02 | 0.15 | 5.47 | 0.65 | 140.9 | 26.6 | 6.7 | 17.7 | 13.2 | 0.38 | 4.6E+05 | 32637 | 71710 | 11634 |
| | SJ103 | 34.83 | 13.79 | 0.05 | 0.25 | 7.72 | 1.26 | 254.1 | 43.7 | 16.6 | 23.9 | 12.0 | 0.69 | 7.7E+05 | 22704 | 36894 | 7058 |
| | SJ101 | 34.76 | 13.79 | 0.06 | 0.30 | 6.83 | 1.45 | 282.9 | 58.4 | 23.6 | 30.6 | 14.5 | 0.77 | 5.4E+05 | 10224 | 51084 | 7807 |
| | SJ108 | 33.69 | 13.06 | 0.11 | 0.29 | 8.31 | 2.55 | 410.2 | 71.2 | 39.5 | 25.4 | 12.2 | 1.56 | 7.7E+05 | 20575 | 55341 | 10759 |
| 19/05/10/ | RV001 | 36.74 | 16.60 | 0.02 | 0.11 | 0.85 | 0.57 | 128.1 | 23.4 | 7.4 | 13.7 | 12.8 | 0.54 | 1.2E+04 | 10320 | 41180 | 80780 |
| | ZI032 | 36.86 | 16.87 | 0.02 | 0.10 | 0.61 | 0.40 | 151.7 | 35.0 | 9.0 | 20.9 | 16.2 | 0.43 | 2.6E+03 | 21300 | 59640 | 45520 |
| | SJ107 | 36.74 | 17.06 | 0.02 | 0.11 | 0.62 | 0.38 | 135.7 | 36.5 | 16.0 | 15.1 | 18.8 | 1.06 | 5.2E+02 | 4260 | 48280 | 24410 |
| | SJ105 | 35.73 | 17.85 | 0.03 | 0.14 | 1.83 | 0.81 | 232.8 | 37.5 | 12.0 | 19.7 | 11.3 | 0.61 | 1.9E+04 | 17040 | 301920 | 173120 |
| | SJ103 | 33.31 | 18.11 | 0.05 | 0.16 | 4.60 | 1.88 | 360.5 | 42.3 | 17.2 | 19.3 | 8.2 | 0.89 | 3.1E+05 | 8520 | 31240 | 28290 |
| | SJ101 | 25.57 | 19.55 | 0.22 | 0.14 | 22.40 | 4.33 | 606.5 | 51.2 | 20.9 | 24.0 | 5.9 | 0.87 | 3.3E+06 | 2840 | 24140 | 23680 |
| | SJ108 | 20.89 | 18.81 | 0.46 | 0.05 | 34.81 | 9.52 | 830.2 | 55.2 | 24.1 | 26.1 | 4.7 | 0.92 | 3.7E+06 | 4260 | 11360 | 12110 |
| 24/[illegible] | RV001 | 35.19 | 21.10 | 0.00 | 0.16 | 1.95 | 0.20 | 93.02 | 22.5 | 6.1 | 14.2 | 16.9 | 0.43 | 2.2E+03 | 7855 | 26961 | 26413 |
| | ZI032 | 34.70 | 21.46 | 0.00 | 0.15 | 2.02 | 0.33 | 151.6 | 39.0 | 10.9 | 23.9 | 18.0 | 0.45 | 9.1E+03 | 10693 | 55721 | 40365 |

| | | | | | | | | | | | | | | | | | |
|---|---|---|---|---|---|---|---|---|---|---|---|---|---|---|---|---|---|
| | SJ107 | 33.18 | 21.99 | 0.02 | 0.13 | 2.56 | 0.63 | 205.8 | 30.4 | 5.9 | 23.1 | 10.3 | 0.25 | 1.5E+04 | 7956 | 120615 | 67228 |
| | SJ105 | 34.57 | 22.23 | 0.00 | 0.11 | 1.77 | 0.40 | 161.8 | 32.9 | 7.4 | 22.9 | 14.2 | 0.32 | 3.6E+03 | 15609 | 59598 | 64778 |
| | SJ103 | 34.15 | 22.09 | 0.04 | 0.15 | 1.95 | 0.64 | 247.4 | 24.3 | 7.7 | 13.7 | 6.9 | 0.56 | 6.7E+03 | 18358 | 80883 | 107599 |
| | SJ101 | 21.35 | 23.07 | 0.37 | 0.34 | 73.69 | 8.91 | 889.1 | 68.4 | 37.1 | 23.6 | 4.5 | 1.57 | 3.4E+06 | 14190 | 177375 | 267915 |
| | SJ108 | 31.86 | 22.61 | 0.04 | 0.38 | 4.00 | 1.48 | 370.6 | 50.5 | 16.7 | 30.6 | 9.5 | 0.55 | 1.9E+05 | 106550 | 233970 | 108210 |
| 15/07/10/ | RV001 | 32.87 | 27.43 | 0.01 | 0.18 | 4.69 | 0.43 | 170.2 | 23.4 | 7.4 | 13.7 | 9.6 | 0.54 | 3.9E+05 | 6250 | 14200 | 8790 |
| | ZI032 | 30.98 | 28.47 | 0.04 | 0.14 | 4.17 | 0.62 | 339.7 | 35.0 | 9.0 | 20.9 | 7.2 | 0.43 | 4.2E+05 | 7100 | 7100 | 19760 |
| | SJ107 | 29.32 | 28.58 | 0.09 | 0.17 | 9.00 | 1.66 | 398.3 | 36.5 | 16.0 | 15.1 | 6.4 | 1.06 | 2.5E+05 | 2130 | 8520 | 7320 |
| | SJ105 | 31.30 | 29.12 | 0.04 | 0.14 | 7.82 | 0.92 | 252.2 | 37.5 | 12.0 | 19.7 | 10.4 | 0.61 | 3.2E+05 | 7100 | 21300 | 11640 |
| | SJ103 | 30.59 | 28.90 | 0.05 | 0.14 | 3.83 | 1.15 | 407.6 | 54.1 | 20.2 | 26.6 | 9.3 | 0.76 | 1.5E+05 | 7810 | 8520 | 6280 |
| | SJ101 | 29.69 | 31.06 | 0.02 | 0.50 | 4.53 | 0.86 | 350 | 60.4 | 13.5 | 41.2 | 12.1 | 0.33 | 5.8E+05 | 5680 | 2840 | 25130 |
| | SJ108 | 30.28 | 29.79 | 0.05 | 0.22 | 3.41 | 0.85 | 400 | 35.5 | 8.5 | 23.6 | 6.2 | 0.36 | 6.4E+05 | 1420 | 21300 | 19260 |
| 19/08/10/ | RV001 | 34.54 | 24.88 | 0.00 | 0.19 | 1.05 | 0.18 | 125.9 | 21.8 | 4.1 | 11.8 | 12.1 | 0.64 | 3.0E+03 | 5870 | 19880 | 3150 |
| | ZI032 | 33.38 | 24.48 | 0.03 | 0.16 | 1.40 | 0.47 | 237.2 | 30.4 | 7.0 | 20.8 | 9.0 | 0.34 | 1.3E+04 | 7100 | 25560 | 2000 |
| | SJ107 | 34.37 | 24.32 | 0.00 | 0.07 | 1.18 | 0.37 | 132.5 | 24.8 | 4.8 | 15.4 | 13.1 | 0.31 | 1.9E+04 | 4260 | 113600 | 2870 |
| | SJ105 | 34.19 | 24.34 | 0.00 | 0.16 | 1.30 | 0.22 | 149.1 | 22.5 | 5.6 | 14.4 | 10.6 | 0.39 | 1.8E+04 | 2840 | 66740 | 2450 |
| | SJ103 | 34.57 | 24.55 | 0.00 | 0.18 | 1.16 | 0.20 | 114.8 | 24.5 | 4.5 | 13.0 | 15.0 | 0.64 | 3.1E+04 | 2840 | 35500 | 4180 |
| | SJ101 | 32.29 | 24.98 | 0.02 | 0.17 | 2.40 | 0.72 | 223.4 | 36.8 | 9.3 | 24.2 | 11.5 | 0.38 | 2.8E+05 | 7100 | 4260 | 4980 |
| | SJ108 | 31.50 | 26.30 | 0.05 | 0.20 | 0.89 | 1.05 | 515.3 | 45.8 | 12.0 | 29.5 | 6.2 | 0.41 | 3.3E+05 | 4257 | 51084 | 6697 |

* Degradation indices

TABLE S2 Eigenvalues of correlation matrix, and related statistics

| Value number | Eigenvalue | % Total variance | Cumulative Eigenvalue | Cumulative % |
|---|---|---|---|---|
| 1 | 4.328521 | 48.09468 | 4.328521 | 48.0947 |
| 2 | 1.976932 | 21.96591 | 6.305454 | 70.0606 |
| 3 | 1.074673 | 11.94081 | 7.380127 | 82.0014 |
| 4 | 0.664392 | 7.38213 | 8.044519 | 89.3835 |
| 5 | 0.509346 | 5.65940 | 8.553865 | 95.0429 |
| 6 | 0.290525 | 3.22806 | 8.844390 | 98.2710 |
| 7 | 0.129794 | 1.44215 | 8.974184 | 99.7132 |
| 8 | 0.025809 | 0.28676 | 8.999993 | 99.9999 |
| 9 | 0.000007 | 0.00008 | 9.000000 | 100.0000 |
